# Language-Model-Based Detection of Genetic Editing in Bacteria

**DOI:** 10.64898/2026.09.17.751846

**Authors:** Edan Gabay, David Burstein

**Affiliations:** The Shmunis School of Biomedicine and Cancer Research, Tel Aviv University, Israel

**Keywords:** Biosecurity, Genomics, Genetic Engineering Detection, Transformers

## Abstract

Recent advances in genome editing allow easy genetic manipulation of bacteria, providing them with new traits, some of which could be hazardous, e.g. enhanced virulence or extended resistance to antibiotics. The ability to detect artificially modified bacteria is crucial for identifying potential bio-threats. However, malicious genome editing could be challenging to trace due to the natural exchange of genes among bacteria through horizontal transfer. After curating extensive datasets including natural genomes and simulated edited genomes, we utilized a natural language processing approach to detect edited genomes. We developed a transformer-encoder-based machine-learning classifier that, instead of analyzing words in sentences, models gene families in genomes. After training the model on our datasets, it is able to accurately detect genes artificially added to bacterial genomes due to their unnatural context. Our approach provides a scalable method for identifying engineered sequences without relying on specific marker genes, with potential applications in biosecurity, agriculture, GMO regulation and more.

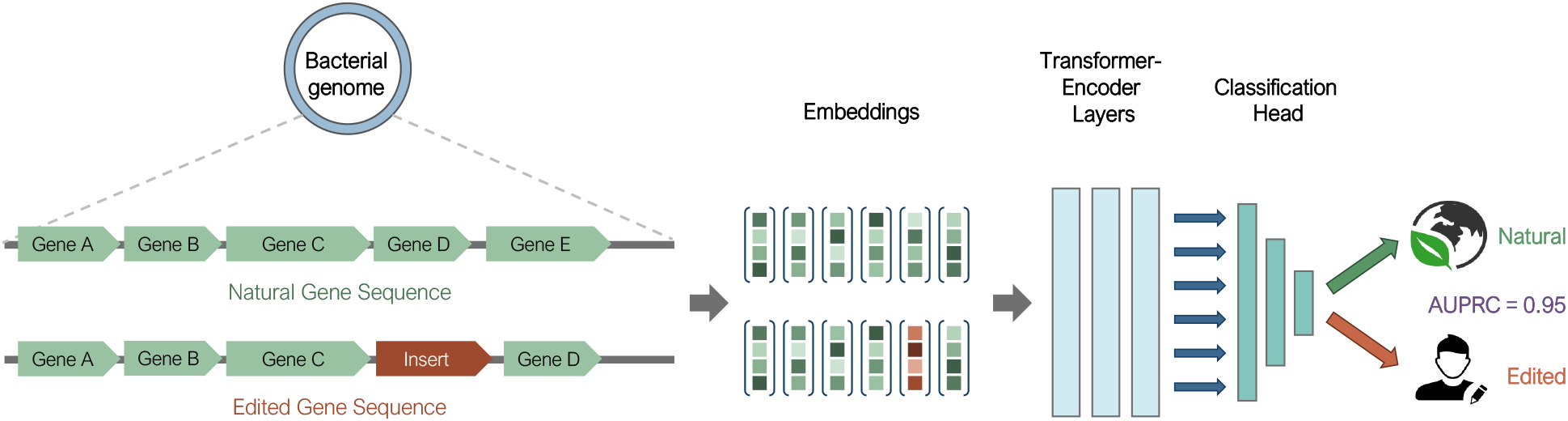

## 1 Introduction

Genome editing involves making precise changes to the DNA of an organism to alter its genetic material, thus changing some of its properties. The recent development of genomic manipulation tools, such as CRISPR-Cas9 editing [1], makes bacterial genome editing extremely affordable and accessible. Such editing is frequently used for biomedical research, agriculture, and various clinical and biotechnological applications [2].

The accessibility and ease of use of advanced genome editing tools could make it straightforward for a malicious user to insert genes that would provide hazardous properties to bacteria, such as enhanced pathogenicity or toxin-producing genes. Further, recent developments in “scarless” genome editing allow removing any selection genes used in the editing process, leaving a genome without clear markers that could enable editing identification [3]. Combined, this calls for the development of tools to identify editing events. In addition to biosecurity-related applications, the ability to identify genetic editing in bacteria has implications across multiple industries and scientific fields. For example, in the food industry and agriculture, reliable detection methods are essential for ensuring compliance with GMO regulations [4]. Medical research and therapeutics rely on these methods to ensure the purity of genetically modified cell lines and monitor therapeutic genes in clinical trials [5, 6]. The development of robust, versatile methods for detecting genetic editing has the potential to address these diverse needs across multiple sectors.

However, detection of genetic editing events is far from being trivial, as bacteria tend to naturally exchange genes via horizontal gene transfer (HGT) through conjugation, transduction, transformation and additional mechanisms [7, 8]. This broad range of possible HGT events complicates the task of distinguishing between HGT and gene insertion through editing. Furthermore, traditional methods of detecting genetic editing events, often reliant on identifying marker genes, may fall short in effectively distinguishing between natural and artificially modified genomes - especially when dealing with scarless editing techniques.

A modern approach for genetic editing identification was developed by Fogel et al. [9]. They trained a neural network to detect genetic engineering in *Bacillus subtilis*, based on *k*-mer distribution differences from alignments to 145 native *B. subtilis* genomes. The model was trained on both natural and simulated engineered *B. subtilis* genomes. However, this model is limited to *B. subtilis*, and is hard to recreate on most other bacteria because it is limited by the availability of genomes. *B. subtilis* is an extremely well-studied model organism with numerous genomes available, and this is not the case for the vast majority of bacteria.

We propose addressing this challenge by leveraging methodologies from the field of Natural Language Processing (NLP), which can be applicable to diverse bacterial genomes. NLP is a field of computational linguistics that aims to analyze and generalize large textual datasets using mathematical and statistical representations. One prominent work in this field was word2vec, developed in 2013 [10, 11]. In word2vec, the trained weights of a shallow neural network are used to produce high-dimension vector representations of the target words, termed “word embeddings”, that were shown to capture word semantics effectively.

One of the most groundbreaking advancements in language model architecture was the introduction of LLMs and the transformer architecture. The transformer is a type of neural network architecture that relies on attention mechanisms to draw global dependencies between the different parts of the input data. The attention mechanism enables the model to weigh the importance of different words in relation to each other when processing text, effectively creating a context-dependent representation of each word while taking into account its relationships with all the other words in the sequence. Transformers have become the cornerstone of many state-of-the-art natural language processing models. In addition to achieving exceptional results on natural language data, transformer-based language models have been applied to biological data producing promising results [12–31]. Most of these model the “protein language” using amino acids as words and protein sequences as sentences, or model nucleotide *k*-mers to capture genomic signatures. These sequence-based language models have successfully predicted various protein properties, including structure, amino acid physicochemical features, protein localization, and more. We and others have recently developed NLP approaches at a different level of abstraction, treating genes as words and genomic regions as sentences [32, 33].

In light of the success of these studies, we chose to address the genetic editing detection challenge by utilizing an NLP approach, leveraging the power of genomic context to detect gene insertion. After exploring several modeling and data-generation strategies, we developed a transformer-encoder-based model, aiming to identify events of gene editing based on the resulting “unnatural” context of genomically edited fragments. Our approach focuses on single-gene insertions, encompassing both arbitrary bacterial genes and genes with potentially malicious functions.

In our simulated benchmarks, the final model achieved strong discrimination between edited and natural sequences, reaching up to 0.95 AUROC and 0.95 AUPRC, with approximately 0.88 test accuracy in the malicious gene insertion setting.

## 2 Methods

### 2.1 Dataset Curation and Simulation

As an initial corpus for the model, we used the one compiled by Miller et al [32]. Briefly, all genomes and metagenomes from NCBI and EBI were acquired and pre-processed in a consistent manner. Pre-processing included gene calling, initial functional annotation, and clustering of genes into families. After filtering out redundancies and short contigs, the dataset contained 360 million genes within their genomic context.

We developed two scalable simulation approaches that are capable of using all available bacterial genomes in order to simulate gene sequences, as detailed below. Briefly, the first method utilizes a curated potentially ‘malicious’ gene database to simulate insertion of these genes into real bacterial genomes, generating simulated edited bacterial sequences. Additionally, we incorporated bacterial sequences containing natural instances of the ‘malicious’ genes as hard-negative examples, alongside random natural bacterial sequences as soft-negative examples. This simulation aims to provide the model with data that enables distinguishing between natural and edited contexts of malicious genes. The second method simulates the insertion of random bacterial genes into unrelated bacterial genome fragments, alongside naturally occurring bacterial sequences as negative samples. This approach results in exceptionally versatile data, enabling a model to discern between natural and unnatural gene context regardless of the type of gene.

In both simulation approaches, samples are represented as lists of consecutive gene identifiers rather than amino acid or nucleotide sequences. This abstraction removes the dependency on specific sequence-level details of the insertion sites and surrounding genomic regions, making the approach highly generalizable. By operating at the gene identifier level, the model can focus on detecting unnatural gene arrangements and contextual patterns without being constrained by the precise molecular characteristics of the insertion sequence and its flanking regions.

### Simulation Strategy: Building Bacterial Genome Datasets

The databases used in our data simulations were compiled as follows:

1. Malicious Gene Insert Simulation:
  a. We curated a set of potentially malicious genes from the following databases:
    i. CARD antibiotic resistance genes database [34]
    ii. DBETH: A Database of Bacterial Exotoxins for Human [35]
    iii. VFDB: Virulence Factor Database [36]
    iv. Bastion: Secreted substrates from five major Gram-negative secretion systems (types I-IV and [37])
    After curating the gene insertion dataset, we randomly split it to a train set (85% of the genes) and a test set (15% of the genes). The goal of the split was to avoid overfit of the model on the specific inserted genes and promote generalization, ensuring performance estimates that are representative of previously unseen insert genes.
  b. Bacterial genome selection: in order to train a model capable of generalizing, we aimed to create a scalable simulation that uses as many bacteria as possible. Initially, we chose to use all available reference genomes of Gram-negative bacteria, as most of the bacterial bio-terrorism agents are Gram-negative.
    i. We chose 11 pathogenic Gram-negative bacteria with clinical relevance for the test set: *Brucella spp*., *Burkholderia mallei, Burkholderia pseudomallei, Chlamydia psittaci, Coxiella burnetii, Francisella tularensis, Rickettsia prowazekii, Rickettsia* *rickettsia, Shigella spp*., *Vibrio cholerae*, and *Yersinia pestis*. All remaining available Gram-negative bacterial genomes were used for the train set, excluding bacteria directly related to the test set bacteria (any species from the *Brucella, Burkholderia, Coxiella, Francisella, Rickettsia, Shigella, Vibrio*, or *Yersinia* genera). The goal of the exclusion was to avoid overfit on the test bacteria, ensuring similar performances on novel, previously unseen bacteria.
    ii. Adding Gram-positive Bacteria: While Gram-positive bacteria are generally less frequently employed as bio-weapons, *Bacillus anthracis* is a significant organism of concern in this regard. We thus created an additional Gram-positive dataset. For the Gram-positive test set, we used *Bacillus anthracis* along with its two pathogenic relatives: *Bacillus cereus* and *Bacillus thuringiensis*. We used all remaining available Gram-positive bacterial genomes for the train set.
    iii. All selected genomes were annotated using prokka [38] to predict gene location on the genome and produce initial functional annotation of the protein products.
    iv. Each protein in the genome was then assigned to a protein family identifier, either from KEGG (Kyoto Encyclopedia of Genes and Genomes, [39]) or a hypothetical protein cluster identifier. This was performed by running a DIAMOND search [40] against a database of protein family representatives that was curated by Miller et al. [32]. This representative database was compiled from all assembled metagenomes and genomes (excluding green plants, fungi, and animals) publicly available on NCBI’s and EBI’s databases. Low-ranking matches (E-value above 10-6) were filtered out. Genes with a significant match to KEGG were assigned the identifier of the highest-ranking KEGG hit. Hits with no KEGG hits received the identifier of the best matching hypothetical family representative, as long as it was below the 10-6 E-value threshold. Proteins with no hit above this threshold were marked with a designated “Unknown” identifier.
    v. Using the assigned identifiers, the contigs (assembled genomic segment) were converted from a DNA sequence to a sequence of identifiers, each representing a protein family.
    vi. The process of assigning protein family identifiers was repeated for each gene in the insert gene databases.
2. Random Bacterial Gene Insert Simulation: We perform step 1b as described above. This simulation strategy did not require a gene insert set, as random bacterial genes from the genome database were inserted instead.

### 2.1.2 Simulation Strategy: Train and Test Set Compilation

1. Malicious Gene Insert Simulation:
  a. As an initial training set, we created 24 million samples composed of simulated insertions and naturally occurring genomic segments, as follows (see Figure 1):
    i. Each sample was a sequence of five protein identifiers.
    ii. 50% of the samples were natural sequences (labeled ‘negative’ for the model). The negative samples were five proteins encoded consecutively on the genome of the chosen bacteria, as follows:
      A. 25% of the samples were ‘hard negatives’: natural sequences from the chosen bacterial genomes that included genes from the insert gene dataset in their natural context. For the simulation, we used all existing hard negative samples in the train bacteria.
      B. 25% of the samples were ‘soft negatives’: random natural sequences from the chosen bacterial genomes.
    iii. 50% of the samples had simulated insertions (labeled ‘positive’ for the model). The positive samples were four proteins encoded consecutively on the genome of the chosen bacteria, with one additional gene from the insert database added randomly into the sequence to create a series of five proteins. The positive samples were created as follows:
      A. 25% of the samples with simulated insertions that used insert genes that also appeared in their natural context in the ‘hard negative’ samples.
      B. 25% of the samples with simulated insertions that used randomly selected genes from the insert gene dataset.
  b. The test set was compiled in the same manner, using the 11 test bacterial genomes and the test insert gene dataset, with the exception of using all available soft negative samples for the test bacteria instead of sub-sampling them.
2. Random Bacterial Gene Insert Simulation:
  a. We created 45 million samples composed of simulated insertions and naturally occurring genomes, as follows:
    i. Each sample was a sequence of five protein identifiers.
    ii. 50% of the samples were natural sequences (labeled ‘negative’ for the model). The negative samples were five proteins encoded consecutively on the genome of the chosen bacteria. We used all possible negative samples from the train bacteria.
    iii. 50% of the samples had simulated insertions (labeled ‘positive’ for the model). The positive samples were four proteins encoded consecutively on the genome of the chosen bacteria, with one additional gene added to a random position in the sequence to create a series of five proteins. The added gene was randomly sampled from a random bacterial genome from the train set bacteria.
    iv. To compile the final dataset, we sampled 15% of the samples created for each bacteria.
  b. The test set was compiled in the same manner, using the 11 test bacterial genomes and the test insert gene dataset.

**Figure 1:**
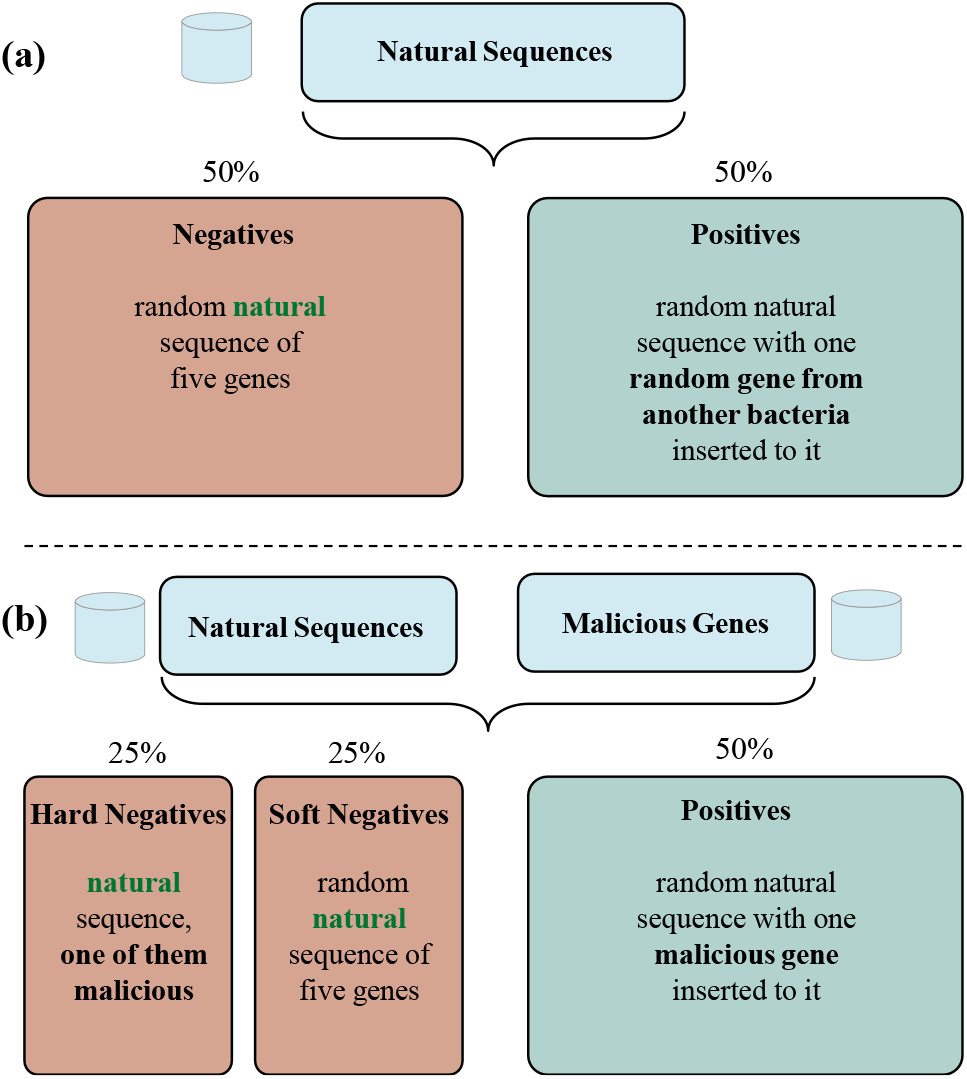
Overview of the gene insertion simulation strategies used for model training. **(a) Random gene insertion simulation**: A negative (natural) sample consists of a naturally occurring sequence of five consecutive genes from a bacterial genome. For the positive set, we sampled a natural sequence of four consecutive genes and inserted an additional, randomly chosen, gene from a different bacterial species to a random position. **(b) Malicious gene insertion simulation**: Positive samples were generated by inserting a gene from a curated dataset of potentially harmful genes (e.g., virulence factors, toxins) to a random position in a natural sequence of four consecutive genes from a bacterial genome. Negative samples consisted of natural sequences of five consecutive genes from a bacterial genome; we sampled both soft negatives (i.e., randomly chosen sequences) and hard negatives (i.e., sequences that naturally contained a gene from the potentially malicious gene database, in its original context). These simulations enabled the model to learn to distinguish between natural genomic contexts and artificially inserted genes.

**Figure 2:**
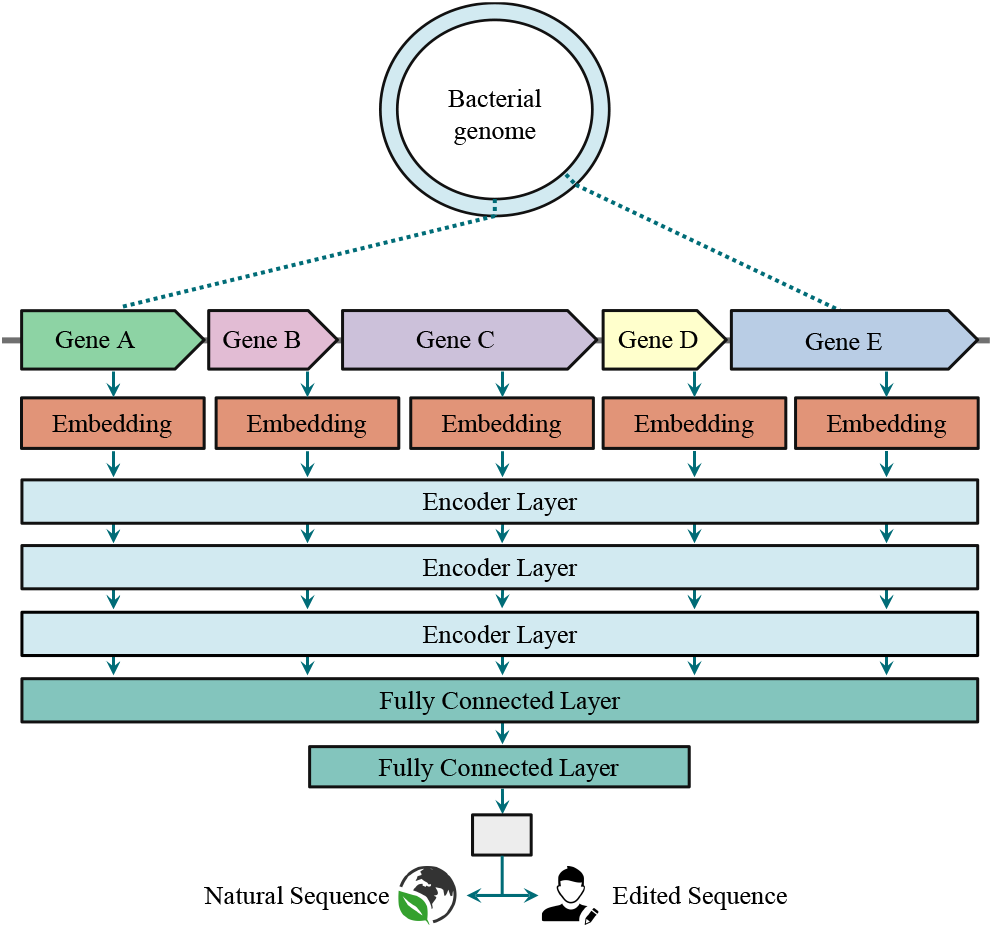
The architecture of the transformer-encoderbased model that was trained on the gene insertion detection task.

### 2.2 Training a Baseline Model

In order to test whether building a machine-learning model for genetic editing detection was feasible, we first trained a simple neural network as a baseline model. For this initial experiment, we only used the Gram-negative bacteria from our datasets for training and testing of the model.

### 2.2.1 Model Input

In a previous study by Miller et al., word2vec was utilized to produce gene embeddings, vectors of 300 values based on their genomic context. We used these embeddings as the initial input for our classification algorithm. In addition to the precomputed embeddings, we added a value marking whether the gene assignment is unknown (0: known, 1: unknown) to distinguish genes that did not have a pre-computed embedding.

### 2.2.2 Baseline Model Architecture and Parameters

We used the Python machine-learning package “PyTorch” [41] to create a simple, five-layered, fully connected neural network for classification. We conducted a hyper-parameter search to determine the most suitable parameters for the model. The parameters explored included the number of encoder layers, dropout rate, learning rate configurations, and the choice between pooling or concatenating model outputs prior to the classifier, as detailed below. The final network included five hidden layers with Rectified Linear Unit (ReLU) as an activation function after each linear layer and a sigmoid activation function for the output layer. The network was trained with the binary cross entropy loss and the “AdamW” optimizer [42].

The model was trained with both simulated datasets mentioned above and achieved good performance (see Results). This encouraged us to develop a more advanced model, better suited for contextual learning: a transformer-encoder-based model.

### 2.3 Training a Transformer-Encoder-Based Model

After proving the feasibility of gene-level context-based models for gene editing detection, we designed and trained a transformer-encoder model. The model was trained in two phases: an initial training phase using the random gene insert simulation, and a fine-tuning phase using the malicious gene insertion simulation.

Instead of randomly initializing a lookup-table for the initial gene embeddings used as the input for the transformer-encoder layers, we used the embeddings trained with word2vec [32], that we had also used as input to the baseline model. These inputs are fed to a series of consecutive transformer-encoder layers. We used the Python machine-learning package “PyTorch” to build the model, and conducted a hyper-parameter search.

### 2.3.1 Model Hyper-parameters

#### Transformer-Encoder Layers

the transformer encoder consisted of three encoder layers, each employing two self-attention heads. To reduce the risk of overfitting during fine-tuning on the malicious gene insertion simulation task, all parameters of the transformer-encoder layers were frozen. The feed-forward sub-network within each encoder layer had a dimensionality of 302, and a dropout rate of 0.1 was applied throughout the model.

#### Classifier Head

we fed the concatenated transformer outputs to the classifier head and received a vector of length (embedding length) × (number of genes). The resulting vector was then fed to a classifier head consisted of a three-layered neural network. We had also tested PyTorch’s ‘AdaptiveMaxPool1d’ layer in order to max-pool the transformer outputs instead of concatenating them, but the concatenation yielded better results. The classifier head used ReLU as an activation function after each linear layer and a sigmoid activation function for the output layer.

### Loss and Optimizer

The network was trained with the binary cross entropy loss and the “AdamW” optimizer.

### 2.3.2. Learning Rate

Our network yielded the best results with a surprisingly low commencing learning rate of 10-4 for the initial training phase and 10-5 for the fine-tuning phase. We used PyTorch’s cosine-annealing learning rate scheduler as a cosine rate decay during each training phase, using a number of iterations equal to the number of total training iterations. We used an eta-min value of 10-6 for the initial training phase and 10-7 for the fine tuning phase. These yielded superior performance in terms of test accuracy, area under ROC and PR curves (AUROC and AUPRC) compared to other learning rate schedulers that were tested.

### 2.4 Training Strategy

Our goal was to train a network that is both versatile in terms of the types of gene inserts it is able to detect, but also likely to succeed in detecting specifically ‘maliciously’ inserted genes. Therefore, we first trained the three-layer transformer-encoder-based model on the random gene insert dataset for ten epochs. Afterwards, we froze the parameters of the transformer-encoder layers and fine-tuned the model on the malicious gene insert dataset. The fine-tuning required 15 epochs to achieve optimal results (The malicious gene insertion dataset is smaller than the random gene insertion dataset). In order to further avoid any overfit or catastrophic forgetting, we used a lower initial learning rate and froze the parameters of the transformer-encoder layers during the fine tuning, as mentioned above.

## 3.1 Results

### 3.1 Scalable Gene Insertion Data simulation

Aiming to develop a highly scalable simulation that includes diverse bacteria and insert genes, two distinct sets of training and testing data were generated. Both datasets represent sequences as lists of gene identifiers, abstracting away sequence-level details to focus on detecting unnatural gene arrangements in bacterial genomes.

The first approach utilizes a curated a database of ‘potentially malicious’ genes to simulate insertion of these genes into real bacterial genomes, thereby simulating ‘maliciously’ edited bacterial sequences. Additionally, we incorporated bacterial sequences containing natural instances of the same ‘potentially malicious’ genes as hard-negative examples, along with randomly selected naturally occurring bacterial sequences as soft-negative examples. This simulation is designed to provide the model with data that can enable differentiation between natural and altered contexts of ‘potentially malicious’ genes. The simulation process includes the following steps (further detailed in the Methods section):

- Selection of insert gene dataset: we selected gene databases of genes for antibiotic resistance (CARD [34]), bacterial exotoxins (DBETH [35]), virulence factors (VFDB [36]) and effectors (BastionHub [37]). To prevent leakage, we split the insert gene dataset into a train set and a test set with distinct sets of genes.
- Bacterial genome selection: We chose to use all available reference genomes from NCBI’s WGS database. We split chosen genomes to train genomes and test genomes to prevent leakage, choosing 11 pathogenic bacteria with clinical relevance for the test genomes, and all available reference genomes of different genera for the training dataset.
- Gene editing was simulated by randomly inserting genes from the chosen databases between two consecutive genes of the selected genome. For the negative sequences, we used naturally occurring sequences from the chosen bacteria. We used a numeric vector representation for each gene (embedding) that was precomputed based on the genomic context of the gene within various genomes and metagenomes [32].
- The second method simulates the insertion of random bacterial genes into unrelated genomic regions supplemented by naturally occurring bacterial sequences as negative samples. This strategy results in a modular database, enabling a model to discern between natural and unnatural gene organizations, irrespective of the type of gene inserted into them. This simulation process includes selecting bacterial genomes in the same manner as in the first simulation. Gene editing was then simulated by inserting a random bacterial gene between two randomly-selected consequitive genes of a different genome. For the negative sequences, we took naturally occurring sequences from the chosen bacteria. Afterwards, we used gene embeddings as in the previous simulation.

### 3.2 Training a Baseline Neural Network Based Model

#### 3.2.1 Baseline Random Gene Insert Model

Using the random gene insertion dataset, we created a versatile dataset of Gram-negative bacteria. The dataset included 45 million samples, half of which are non-edited (negative) and half edited (positive). The data was used to train a fully-connected neural network classifier to distinguish between edited and naturally occurring bacterial gene sequences.

This classification task is particularly challenging, since the model must distinguish between natural and edited gene sequences based solely on their contextual patterns, without any information about the gene function. Despite the difficulty of the task, this baseline model still achieved reassuring results: an AUROC value of 0.84, an AUPRC value of 0.79 and a test accuracy of 0.75 (for a threshold of 0.5, see Table 1).

**Table 1:**
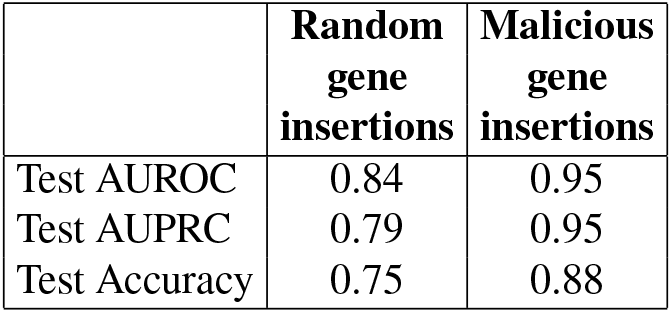
Baseline model performance on two simulation approaches: the malicious gene insertion simulation and the random gene insertion simulation. Test accuracy was calculated for a threshold of 0.5.

|  | **Random**<br>**gene**<br>**insertions** | **Malicious**<br>**gene**<br>**insertions** |
| --- | --- | --- |
| Test AUROC | 0.84 | 0.95 |
| Test AUPRC | 0.79 | 0.95 |
| Test Accuracy | 0.75 | 0.88 |

#### 3.2.2 Baseline Malicious Gene Insert Model

Using the malicious gene insert simulation method, we created an extensive dataset of Gram-negative bacteria containing 24 million samples, half of which are non-edited (negative) and half edited (positive). The negative samples included both ‘hard negative’ samples, which are naturally occurring sequences that include genes from the insert gene dataset in their natural context, as well as ‘soft negative’ examples, which are randomly selected naturally occurring gene sequences. The data was used to train a fully connected neural network classifier aiming to distinguish between naturally occurring bacterial gene sequences and genomes supplemented with a potentially malicious gene. The model performed very well on this dataset, with both the area under the ROC curve (AUROC) and the area under the PR curve (AUPRC) achieving a value of 0.95, and a test accuracy of 0.88 (for a threshold of 0.5).

We evaluated three distinct negative sampling strategies for training the model: using exclusively soft negatives to train the model, using exclusively hard negatives to train the model, and using a balanced combination of both soft and hard negatives in equal proportions.

The results demonstrated that incorporating both types of negative samples yielded superior performance across all metrics (see Table 2 and Figure 3).

**Table 2:** Performance metrics for the baseline model on the malicious gene insertion approach, across different negative sampling strategies.

|  | **Soft**<br>**Negatives** | **Hard**<br>**Negatives** | **Soft +**<br>**Hard**<br>**Negatives** |
| --- | --- | --- | --- |
| Test AUROC | 0.84 | 0.61 | 0.95 |
| Test AUPRC | 0.85 | 0.60 | 0.95 |
| Test Accuracy | 0.75 | 0.60 | 0.88 |

**Figure 3:**
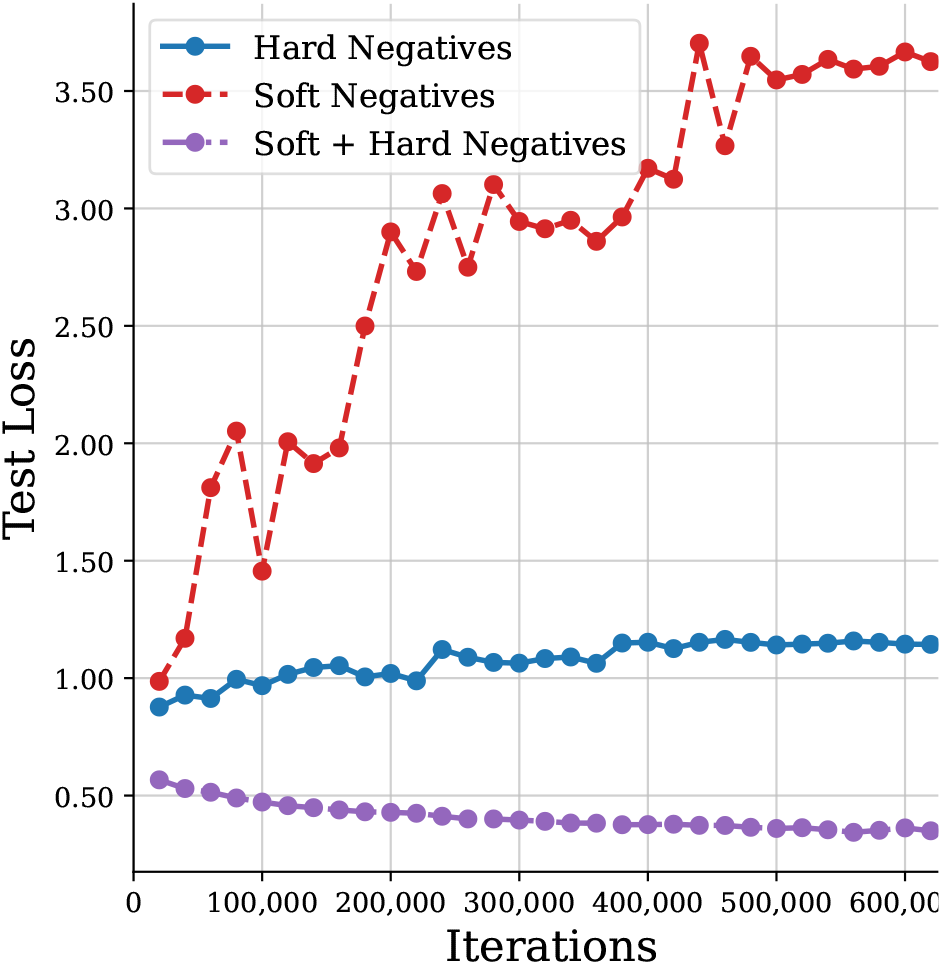
Test loss when training the baseline model on datasets created with the malicious gene insertion approach using different negative sampling techniques.

### 3.3 Training the Transformer-Encoder Model on the Random Gene Insert Simulation Dataset

Using the random gene insert simulation method, we created an extremely versatile dataset with 82 million samples, half of which are non-edited (negative) and half that simulate gene insertion of over 20 million different genes of various functions (positive). The data was used to train a transformer-encoder-based classifier to distinguish between edited and naturally occurring bacterial gene sequences. When trained on a combined Gram-negative and Gram-positive dataset, and using hyper-parameters selected via a dedicated hyper-parameter search, the model received an AUROC of 0.92, an AUPRC of 0.91, and a test accuracy of 0.83 (for a threshold of 0.5). When restricted to Gram-negative bacteria only, performance remained similar, with an AUROC of 0.90, an AUPRC of 0.90, and a test accuracy of 0.81 at the same threshold (see Table 3).

**Table 3:** Performance of the pre-trained model on the task of detecting insertion of different gene sets into Gram-negative bacteria.

| Test Metric | Random Gene Insertion Test Set | Malicious Gene Insertion Test Set |
| --- | --- | --- |
| AUROC | 0.90 | 0.91 |
| AUPRC | 0.90 | 0.91 |
| Accuracy | 0.81 | 0.81 |
| F1 | 0.81 | 0.81 |
| Precision | 0.80 | 0.83 |
| Recall | 0.82 | 0.78 |

#### 3.3.1 Evaluating the Pre-Trained Model on the Malicious Gene Insertion Dataset

The initially trained transformer-encoder model was trained solely on the random gene insertion simulation. To assess whether the model generalizes well to specifically identify potentially malicious gene insertions, we evaluated its performance on this task without any additional training. The results indicate that the initially trained model maintained its predictive capability, achieving nearly identical performance metrics on the malicious gene insertion test set compared to the random gene insertion test set (see Table 3).

### 3.4 Fine-Tuning the Transformer-Encoder-Based Model on the Malicious Gene Insertion Dataset

After training the network on the random gene insert simulation, we froze the parameters of the transformer-encoder layers and fine-tuned the model on the malicious gene insert simulation. Using the first simulation method, we created an extensive dataset with 27 million samples, half of which are non-edited (negative) and half edited (positive). The negative samples included both ‘hard negative’ samples, which are naturally occurring sequences that include genes from the insert gene database in their natural context, as well as ‘soft negative’ examples, i.e., randomly selected naturally occurring gene sequences. On this data, a transformer-encoder-based classifier was trained to distinguish between edited and naturally occurring bacterial gene sequences. The model achieved improved performance with both an AUROC and an AUPRC of 0.95, and a test accuracy of 0.875 (for a threshold of 0.45, see Figure 4 and Table 4).

**Table 4:**
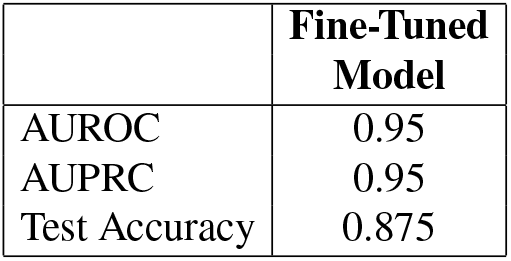
Performance metrics for the transformer-encoder-based model after fine-tuning on the malicious gene insertion dataset.

|  | Fine-Tuned Model |
| --- | --- |
| AUROC | 0.95 |
| AUPRC | 0.95 |
| Test Accuracy | 0.875 |

**Figure 4:**
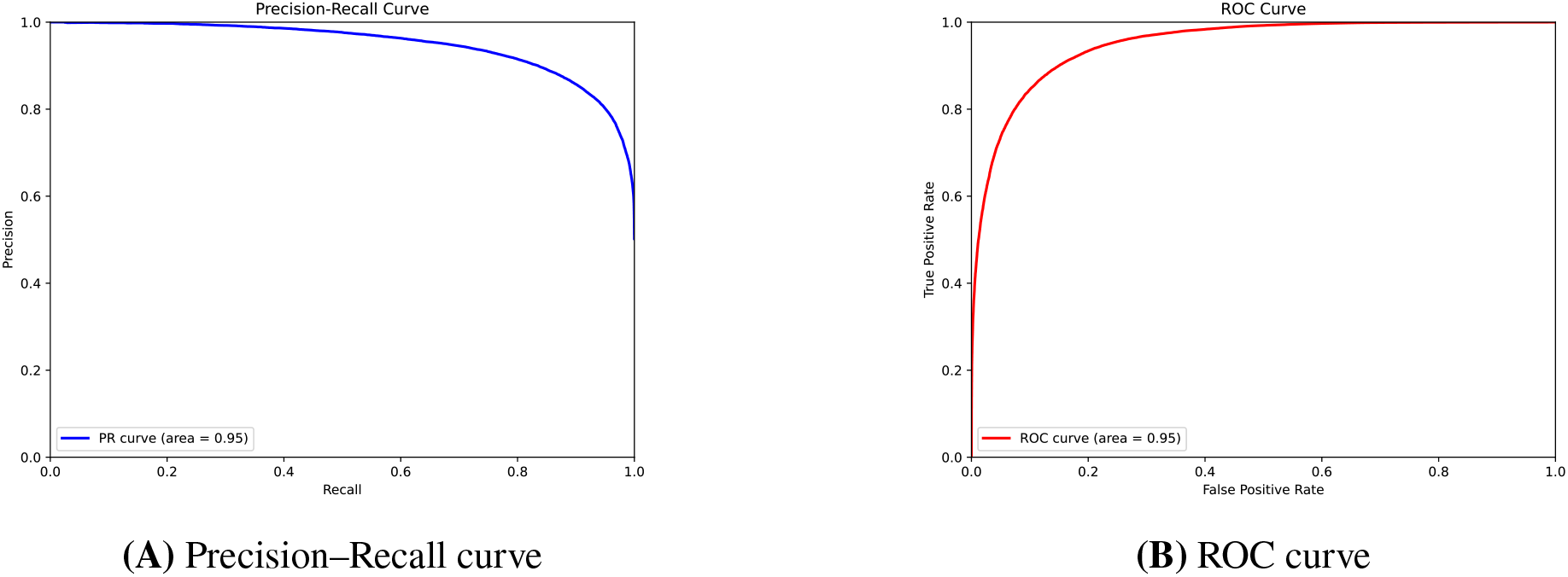
Results for the transformer-encoder-based model trained on the random gene insert simulation (Gram-negative and Gram-positive bacteria), fine-tuned on the malicious gene insert simulation dataset (Gram-negative and Gram-positive bacteria): (A) Precision-Recall curve. (B) ROC curve.

#### 3.4.1 Evaluating the Fine-Tuned Model on the Random Gene Insertion Dataset

To ensure that fine-tuning on the malicious gene insertion simulation did not cause the model to lose its ability to detect general gene insertions, we tested the fine-tuned model on the random gene insertion test set. The model retained most of its predictive power, indicating the advantage of using the two-step training to retain generalization (see Table 5).

**Table 5:** Performance comparison of the initially trained and fine-tuned models on the random gene insertion test set.

| Test Metric | Initially Trained Model | Fine-Tuned Model |
| --- | --- | --- |
| Accuracy | 0.81 | 0.76 |
| F1 | 0.81 | 0.70 |
| AUPRC | 0.90 | 0.84 |
| AUROC | 0.90 | 0.86 |

## 4 Discussion

This study demonstrates the potential of using natural language processing techniques, specifically transformer-encoder-based models, for detecting genome editing events in bacteria. In recent years, transformer-encoder-based classification models have achieved impressive results on various sequence-level prediction tasks, and we found this also applies for gene-level language models classifying natural vs. edited genomes. Our approach leverages the contextual information of genes within genomes to identify artificially inserted sequences. By treating genes as words and genomic regions as sentences, this work leverages language modeling techniques to capture the complex relationships and contexts within bacterial genomes.

We developed two scalable simulation approaches for generating training data, addressing the lack of large-scale databases of genetically edited bacterial genomes. These approaches include a malicious gene insert simulation utilizing curated databases of potentially harmful genes and a random bacterial gene insert simulation for more generalized detection. Both simulation approaches present a sample as a list of consecutive gene-identifiers. Our choice to represent bacterial sequences as gene identifiers rather than nucleotide or amino acid sequences allows the model to focus on broader genomic organization patterns while remaining robust to variations in exact insertion sites and local sequence contexts. This abstraction level is valuable for identifying gene editing occurrences regardless of the precise sequence.

Our transformer-encoder-based model was successfully applied to the task of detecting artificial gene insertion in the simulated datasets. The final model achieved an AUROC and AUPR of 0.95, with a test accuracy of 0.865 on the malicious gene insert dataset. Initial training on the random gene insert dataset followed by fine-tuning on the malicious gene dataset improved the model’s performance and generalization capabilities. Our model demonstrated that genomic context could be effectively utilized to distinguish between natural and artificially inserted gene sequences, even without relying on specific marker genes or other traditional indicators of genetic modification. The resulting model is versatile, capable of detecting both random gene insertions and specifically targeted gene insertions, enhancing its potential for real-world biosecurity applications.

The main limitation of this study is that the model was trained using simulated datasets rather than real-world examples of genetically engineered bacteria. The simulation-based approach was necessitated by the current lack of comprehensive, publicly available datasets containing genetically engineered bacterial genomes. However, the model shows promise for generalization to real-world applications due to its gene-identifier level representation approach. As authenticated samples of genetically modified organisms become available, future work could focus on fine-tuning and testing the model using such datasets. In addition, our model could be extended to detect other types of genetic modifications beyond single gene insertions, such as the insertion of entire operons, non-coding regions and promotor manipulation. Incorporating additional genomic features or metadata could further improve detection accuracy.

In conclusion, this work presents a novel approach for detecting genome editing events in bacteria using transformer-encoder models and gene-level sequence representation. By successfully distinguishing between natural and artificially modified sequences in our simulated datasets, the model demonstrates the potential of applying natural language processing techniques to biosecurity challenges. While further validation with real-world samples is needed, the gene-identifier level abstraction and strong performance metrics suggest promising applications in screening for potentially harmful genetic modifications. As genome editing technologies continue to advance, scalable computational approaches will become increasingly valuable for monitoring and detecting potential misuse of synthetic biology.

## Acknowledgments

This work was supported in part by the Israel Science Foundation [grant number 355/23] and by a fellowship to EG from the Edmond J. Safra center for Bioinformatics at Tel Aviv University. We thank Ella Ranon for critical reading of the manuscript.

